## Supplementary material for "Non-hydrolyzable acetyllysine analogs to study protein acetylation in vitro and in cells": SupInfo

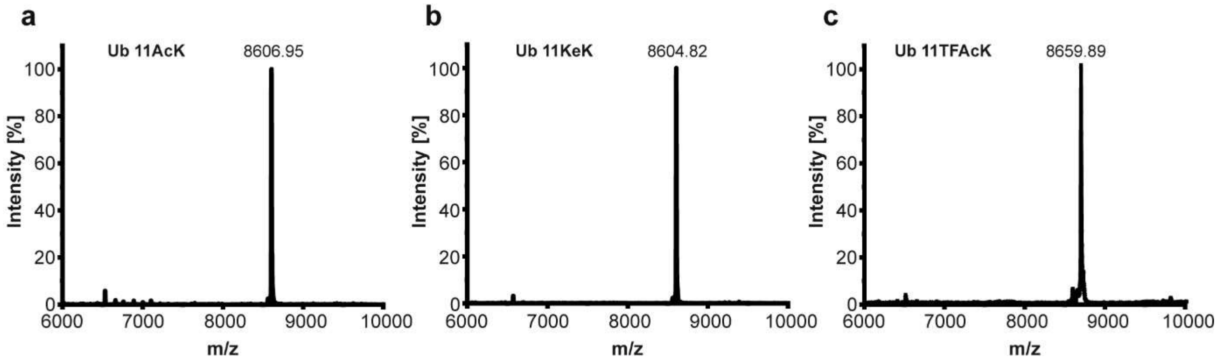


**Supplementary Fig. 1. ESI-MS spectra of site-specifically acetylated Ub variants.** ESI-MS analysis demonstrates the quantitative incorporation of **a** acetyllysine (AcK), **b** ketolysine (KeK), and **c** trifluoroacetyllysine (TFAcK) into Ub at position 11. The calculated masses of Ub 11AcK, 11KeK, and 11TFAcK are 8606.84 Da, 8605.82 Da, and 8660.74 Da, respectively.

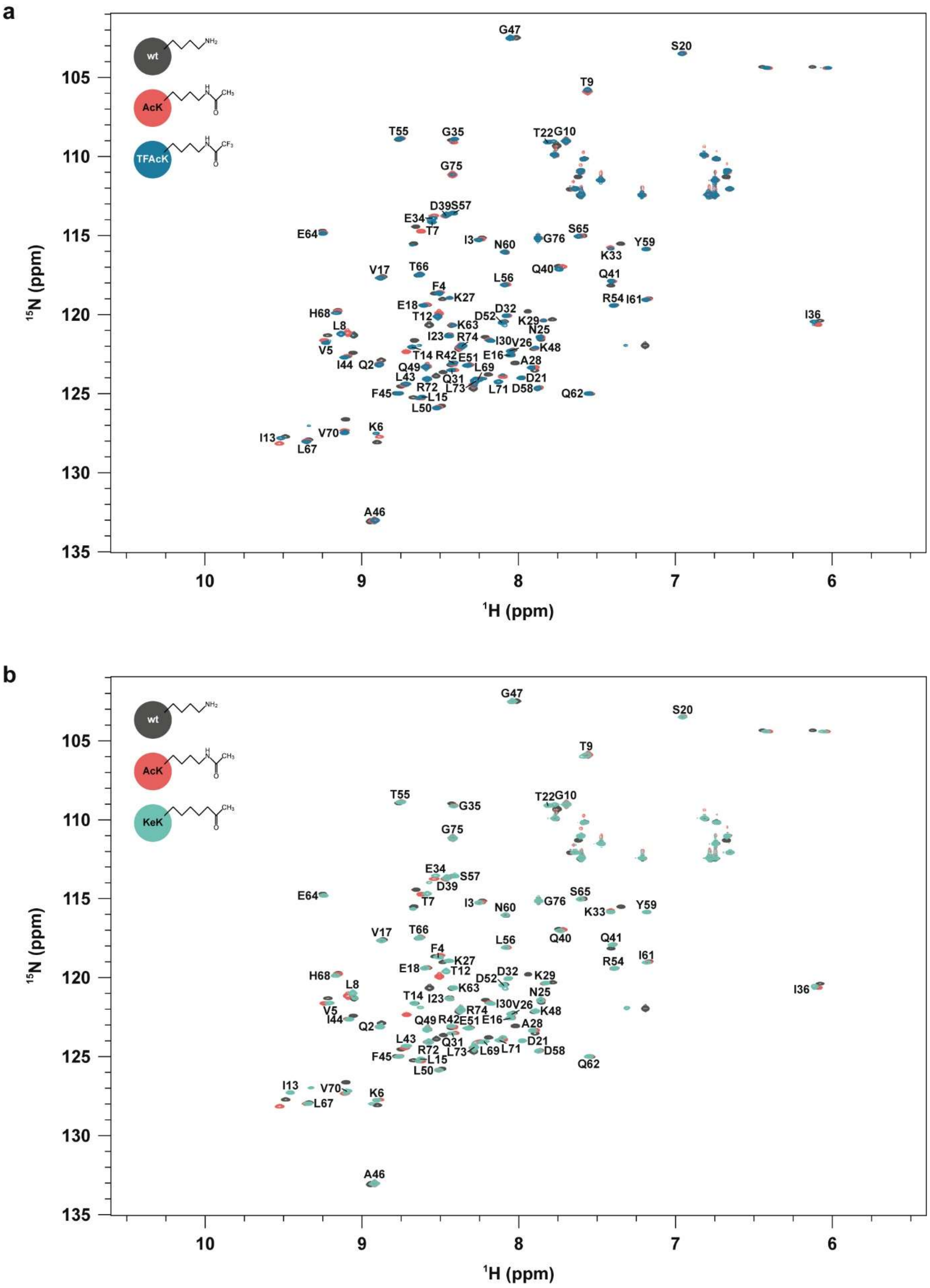


**Supplementary Fig. 2. NMR spectra of acetylated Ub K11 variants. a-b,** Two-dimensional $^1$H-$^{15}$N HSQC NMR spectra of **a** Ub 11TFAcK (TFAcK) and **b** Ub 11KeK (KeK). The corresponding CPSs of the Ub variants are shown in Fig. 2. Resonance signals originating from Ub 11TFAcK and Ub 11KeK are colored in blue and pale green, respectively, and are labeled by using the one letter code. A superimposition with the spectrum of unmodified Ub (wt) colored in anthracite and Ub 11AcK (AcK) colored in red is provided in **a** and **b.** Note that the respective backbone amide nitrogen of the modified lysine residues is unlabeled resulting in no cross peak in the respective spectrum.

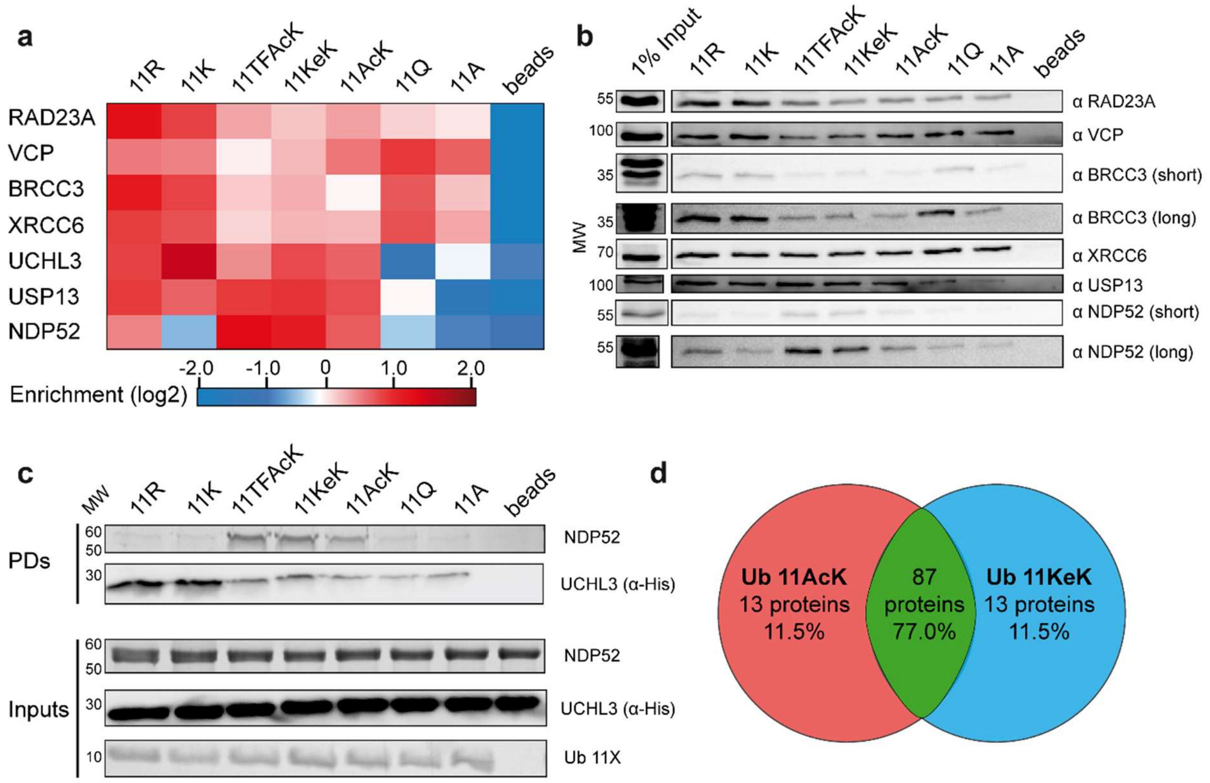


**Supplementary Fig. 3. Selected interactors show a similar binding behavior to Ub 11 variants in different experimental set-ups. a,** Selected interactors of Ub 11 variants with a distinct binding pattern. Red indicates enrichment, whereas blue indicates lack of enrichment. **b,** Confirmation of interactions by western blot analysis. 20% of the elution fraction of an affinity enrichment experiment were subjected to western blot analysis with specific antibodies against the proteins indicated. Input represents 1% of the HEK293T cell extract used for affinity enrichment. **c,** Verification of interactions by *in vitro* binding assays using recombinant NDP52 and UCHL3 and the Ub variants indicated. Binding reactions (PDs) were analyzed by SDS-PAGE followed by Coomassie blue staining for NDP52 and western blot analysis for UCHL3. Inputs, 10% of the proteins used in the binding assay were subjected to SDS-PAGE followed by Coomassie blue staining. For His6-UCHL3 detection an anti-His antibody coupled to HRP was used (α-His). Running positions of molecular mass markers (MW) are indicated in **b** and **c**. In **b**, the antibodies used as well as exposure times (short or long) are indicated. **d**, Venn diagram of top 100 significantly enriched interactors for Ub 11AcK (red) and Ub 11KeK (blue). Number and percentage of unique proteins and the overlap between Ub11Ac and 11KeK (intersection, green) are indicated.

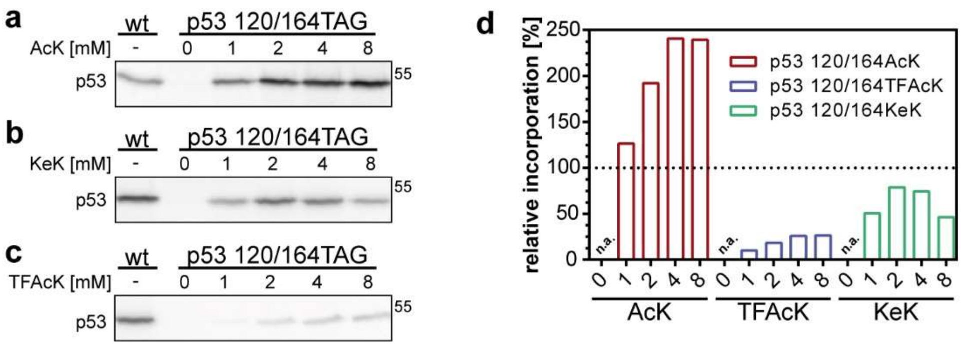


**Supplementary Fig. 4. Generation of double acetylated p53 variants in H1299 cells.** H1299 cells were transfected with expression constructs encoding wild-type p53 (wt) or p53 with TAG stop codons at position 120 and 164 and the AcK-RS/tRNA pair as described in Methods. Growth medium was supplemented with increasing concentrations of **a** AcK, **b** KeK, and **c** TFAcK as indicated. 24 h upon transfection, p53 levels were determined by western blot analysis using the p53-specific antibody DO-1. **d,** Quantification of relative p53 levels in the presence of AcK, TFAcK, and KeK. Levels of wild-type p53 (wt) were set to 100 percent.

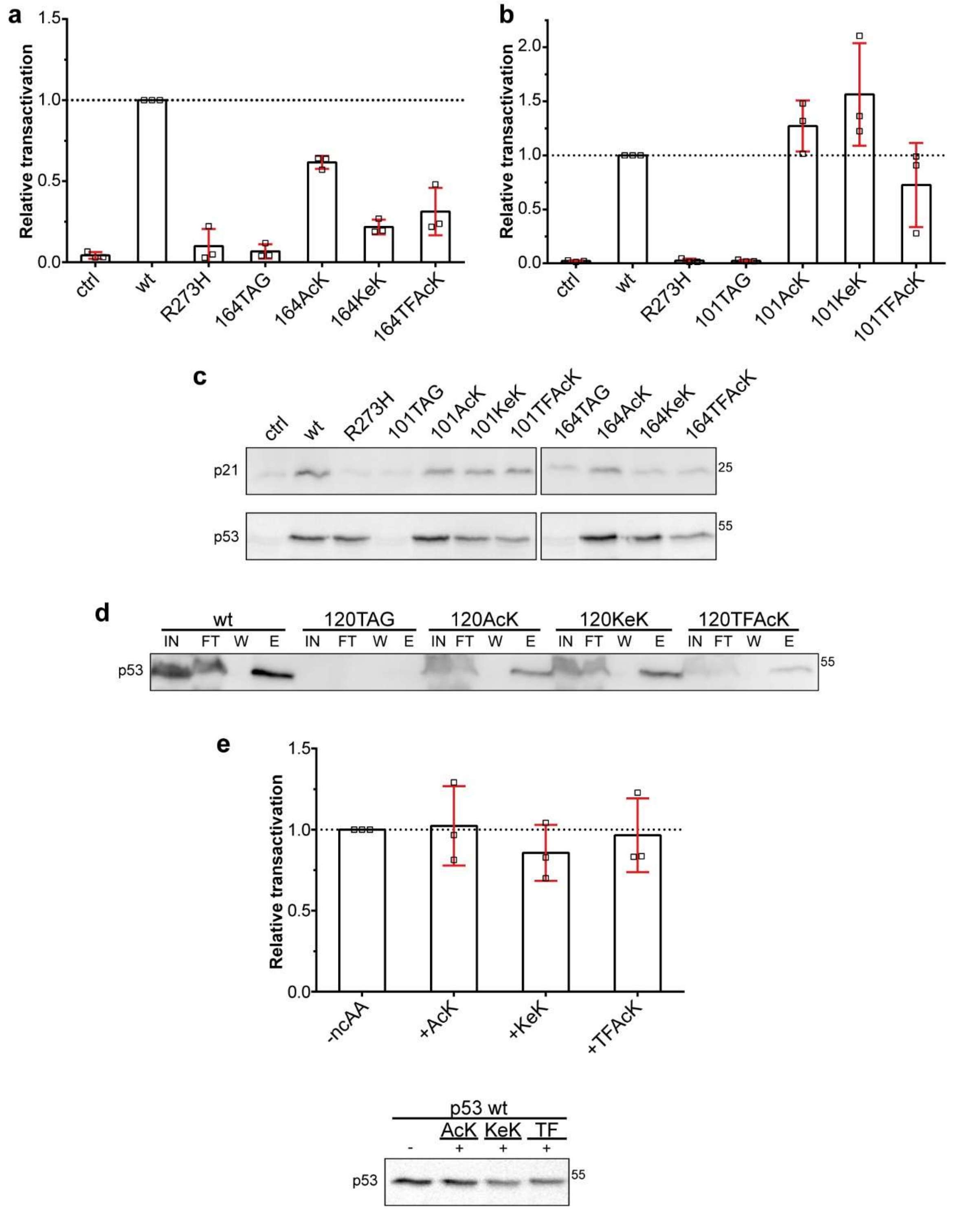


**Supplementary Fig. 5. a-b,** H1299 cells were transfected with a p53-responsive reporter construct encoding luciferase (containing the p21 response element for p53) in the absence (ctrl) or presence of constructs encoding wild-type p53 (wt), the tumor-derived p53 mutant R273H, p53 with a TAG stop codon at position 164 (**a**) or position 101 (**b**), and the AcK-RS/tRNA pair. In addition, cells transfected with the p53-164TAG or p53-101TAG construct were grown in the absence or presence of AcK (164AcK, 101AcK), KeK (164KeK, 101KeK), or TFAcK (164TFAcK, 101TFAcK). 24 h upon transfection, luciferase activity was determined as a measure for p53 activity and the relative values obtained were adjusted for transfection efficiency (with n = 3 ± SD independent experiments). **c** H1299 cells were transfected as in **a**, **b** in the absence of a p53-responsive reporter construct. 24 h upon transfection, levels of endogenous p21 protein and of ectopically expressed p53 were determined by western blot analysis using a p21-specific antibody and the p53-specific DO-1 antibody, respectively. **d** Affinity enrichment of p53 variants for subsequent PRM analysis (Fig. 5c). H1299 cells were transfected with expression constructs for C-terminally Strep-II tagged wild-type p53 (wt) or p53-120TAG in the absence (120TAG) or presence of AcK (120AcK), KeK (120KeK), and TFAcK

(120TFAcK) as indicated. 24 h upon transfection, cell extracts were prepared and p53 variants enriched via Strep-Tactin® beads. Enrichment was verified by western blot analysis using the p53-specific antibody DO-1. IN, Input; FT, flow through; W, wash; E: elution. 1% of the total of the respective fractions were analyzed. **e** Presence of AcK, KeK, or TFAcK does not affect the transactivation capacity of wild-type p53. H1299 cells were transfected with a p53-responsive expression construct encoding luciferase (containing the p21 response element for p53) and a construct encoding wild-type p53 in the absence (-ncAA) or presence of AcK, KeK, TFAcK (TF) as indicated. 24 h upon transfection, luciferase activity was determined as a measure for p53 activity and the relative values obtained were adjusted for transfection efficiency (n = 3 ± SD independent experiments). 20 percent of the extracts were subjected to western blot analysis using the p53-specific DO-1 antibody.

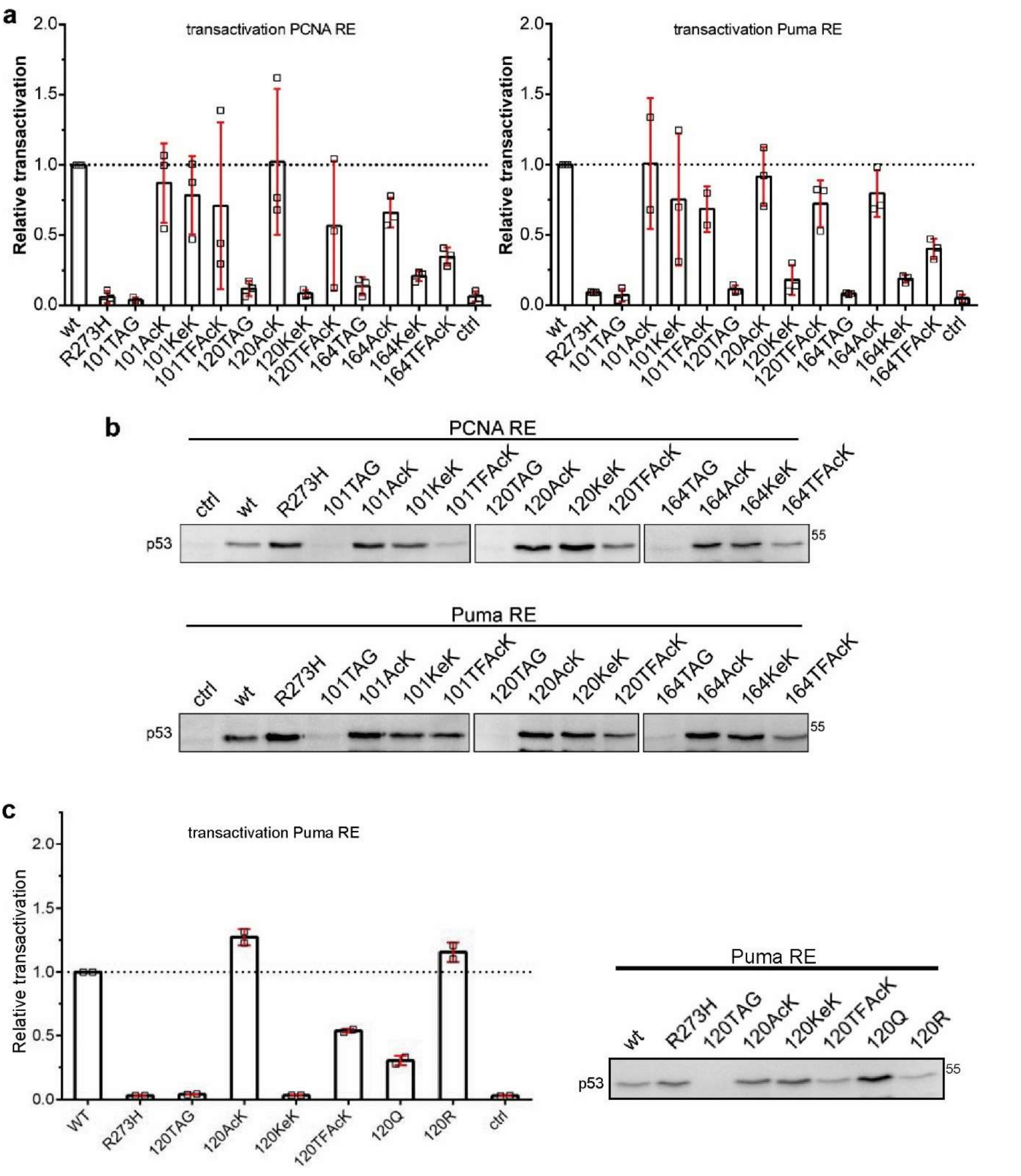


**Supplementary Fig. 6. Transactivation of additional p53-responsive reporters. a,** H1299 cells were transfected with reporter constructs encoding luciferase containing a response element for p53 of either the *PCNA* gene (PCNA RE) or the *PUMA* gene (Puma RE) in the absence (ctrl) or presence of constructs encoding wild-type p53 (wt), the tumor-derived p53 mutant R273H, p53 with a TAG stop codon at position 101, 120, or 164, and the AcK-RS/tRNA pair. In addition, cells transfected with the p53-101TAG, p53-120TAG and p53-164TAG constructs were grown in the absence (X TAG, with X denoting the respective position of the TAG codon) or presence of AcK (X AcK), TFAcK (X TFAcK) or KeK (X KeK). 24 h upon transfection, luciferase activity was determined as a measure for p53 activity and the relative values obtained were adjusted for transfection efficiency (n = 3 ± SD independent experiments. **b,** 20 percent of the extracts were subjected to western blot analysis using the p53-specific DO-1 antibody. **c,** H1299 cells were transfected with the Puma RE as in a. Left panel, Determination of transactivation activity as in a. In addition to wt p53, R273H, p53-120AcK, p53-120KeK, and p53-120TFAcK, the transactivation activity of the p53 variants 120Q and 120R was determined (n = 2 ± SD independent experiments). Right panel, 20 percent of the extracts were subjected to western blot analysis using the p53-specific DO-1 antibody.
